# Environmental modification via a quorum sensing molecule influences the social landscape of siderophore production

**DOI:** 10.1101/053918

**Authors:** Roman Popat, Freya Harrison, Ana C. da Silva, Scott A. S. Easton, Luke McNally, Paul Williams, Stephen P. Diggle

## Abstract

Bacteria produce a wide variety of exoproducts that favourably modify their environment and increase their fitness. These are often termed ‘public goods’ because they are costly for individuals to produce and can be exploited by non-producers (‘cheats’). The outcome of conflict over public goods is dependent upon the prevailing environment and the phenotype of the individuals in competition. Many bacterial species use quorum sensing (QS) signalling molecules to regulate the production of public goods. QS therefore determines the cooperative phenotype of individuals, and influences conflict over public goods. In addition to their regulatory functions, many QS molecules have additional properties that directly modify the prevailing environment. This leads to the possibility that QS molecules could influence conflict over public goods indirectly through non-signalling effects, and the impact of this on social competition has not previously been explored. The *Pseudomonas aeruginosa* QS signal molecule PQS is a powerful chelator of iron which can cause an iron starvation response. Here we show that PQS stimulates a concentration-dependent increase in the cooperative production of iron scavenging siderophores, resulting in an increase in the relative fitness of non-producing siderophore cheats. This is likely due to an increased cost of siderophore output by producing cells and a concurrent increase in the shared benefits, which accrue to both producers and cheats. Although PQS can be a beneficial signalling molecule for *P. aeruginosa*, our data suggests that it can also render a siderophore-producing population vulnerable to competition from cheating strains. More generally our results indicate that the production of one social trait can indirectly affect the costs and benefits of another social trait.

## 1. Introduction

Bacterial cells secrete numerous extracellular factors to favourably modify their environment. These include hydrolytic enzymes, protective polymeric matrices for biofilm formation, and biosurfactants that aid motility. The benefits of such exoproducts can accrue both to the producing cell and to neighbouring cells and are therefore termed ‘public goods’ [1]. Public goods are costly for individual cells to produce, and cooperating populations are consequently at risk of social exploitation by non-producing ‘cheats’ [1,2]. In theory, cheats can outcompete cooperators because they do not incur the cost of public goods production, but derive benefits from the cooperation of others. Whether cooperation persists over evolutionary time in the face of the advantages of cheating is largely dependent on aspects of population structure that act to align individual interests [3].

Many cooperative behaviours seen in bacteria are regulated at the population level by cell-to-cell communication or quorum sensing (QS) systems [4–6]. Cells produce and release QS molecules to regulate the production of a range of public goods which aid in scavenging for nutrients, providing scaffolding for biofilms and facilitating motility. Because these cooperative secretions can be key determinants of successful growth or persistence, there has been considerable interest in the impact of QS on ecological competition between different genotypes or strains of bacteria [7–9]. For example, mutant genotypes which do not respond to QS molecules, and consequently produce fewer or no public goods (even though the loci that directly encode these public goods are intact), have been shown to act as social cheats both *in vitro, in vivo* and in biofilms [8,10–13]. In addition to regulating public goods production, QS molecules have been shown to have non-signalling effects such as immune modulation, cytotoxicity, redox potential and iron binding [14,15]. The impact of these indirect effects by QS molecules on social competition has not previously been explored, and so here we empirically demonstrate how production of a QS molecule can alter the social landscape of a seemingly unrelated trait, siderophore production.

*Pseudomonas aeruginosa* is a Gram-negative opportunistic pathogen which employs a multilayered QS system to regulate a number of public goods, many of which are important for virulence [4,5]. One well-defined *P. aeruginosa* QS signal is the Pseudomonas Quinolone Signal (PQS) [16]. PQS is a member of the 2-alkyl-4(1*H*)-quinolone family of molecules and acts as a QS molecule in the classical sense, in that it interacts with a specific receptor protein, and sets in motion a regulatory cascade leading to increased production of toxins and biofilms [16–18]. PQS also has other biological properties which are distinct from signalling: these include balancing redox reactions, aiding in competition with other species and interacting with cell membranes [17,19,20]. In addition, PQS has iron chelating activity, though it does not act as a true siderophore because it does not directly ferry iron into the cell [21,22]. It has therefore been suggested that PQS may act as an iron trap, aiding in the sequestration, but not in the membrane transport of iron [22].

Moving iron from either a host or the environment into the cell is often achieved by the production of dedicated iron scavenging molecules known as siderophores [23]. *P. aeruginosa* produces two major siderophores, pyoverdine and pyochelin. Pyoverdine has been experimentally demonstrated to be a public good [24] which is exploitable by cheats both *in vitro* and *in vivo* [25,26]. Here, we test whether the iron chelating properties of PQS can change the social landscape of siderophore production. We show that PQS (a) increases the production of pyoverdine and pyochelin, and consequentially decreases the fitness of siderophore producers and (b) increases the relative fitness of siderophore cheats in co-culture with a producing strain. Our findings highlight how direct modification of the environment by one bacterial exoproduct, in this case a QS signal molecule, can indirectly affect the evolutionary dynamics of another social trait.

## 2. METHODS

### (a) Growth media

For a rich, iron-replete growth environment, we used Lysogeny Broth (LB) (10g L^−1^ tryptone, 5g L^−1^ yeast extract, 10g l^−1^ NaCL), and for an iron-limited growth environment we used Casamino Acid (CAA) medium (5g L^−1^ Casamino acids, 1.18g L^−1^ K2HPO_4_.3H_2_0, 0.25gL^−1^ MgSO4.7H_2_0). We prepared both media in dH_2_O and supplemented CAA medium with sodium bicarbonate solution to a total of 20 mM. For all experiments, we inoculated single colonies of the relevant bacterial strain into 5 ml LB and incubated at 37°C at 200 rpm for 18 h. We then washed pre-cultures in the appropriate medium, corrected to an optical density of OD_600_ = 1.0, and inoculated experimental cultures to an initial density of OD_600_ = 0.01.

### (b) Pyoverdine and pyochelin public goods production in response to PQS and HHQ

To study the effects on siderophores of varying concentrations of iron, PQS and its precursor HHQ, we used the strain PAO1Δ*pqsAH*, which is defective in 2-alkyl-4(1*H*)-quinolone production [22], To test whether investment in siderophores increased with added PQS or HHQ, we inoculated a washed pre-culture of PAO1Δ*pqsAH* into 750 *μ*l LB medium containing varying concentrations of PQS and HHQ in microtiter plates and incubated at 37°C for 14 h. Following incubation, we measured the OD_600_ of resulting cultures, and then filtered the cell supernatants. We measured pyoverdine and pyochelin using excitation/emission assays (ex/em wavelengths 400nm/460nm for pyoverdine and 350nm/430nm for pyochelin [27] using a Tecan Multimode plate reader. We corrected fluorescence values by subtracting the fluorescence of a sterile medium blank and assuming a 5% leakage from the pyoverdine into the pyochelin channel, as previously described [27]. We estimated per-cell siderophore production as fluorescence (RFU - Relative Fluorescence Units) divided by culture density (OD_600_).

### (c) Relative fitness of a pyoverdine non-producer

To measure the growth (fitness) of monocultures, we inoculated a washed pre-culture of either PAO1 or PAO1Δ*pvdD/pchEF* into 750 *μ*l CAA medium supplemented with 20 mM NaHCO_3_ containing no addition, or supplementation with either 50 *μ*M PQS, or 100 *μ*g/ml (1.25mM) transferrin in 48 well microtiter plates and incubated at 37°C for 14 h. To study the effect of PQS on the competition between siderophore producers and non-producers we used wild type PAO1 and a mutant that was defective in pyoverdine and pyochelin production and labelled with a constitutive luminescence marker (PAO1Δ*pvdD/pchEF* CTX*lux*). For competition assays, we pre-cultured, washed and density corrected both strains and mixed them to a ratio of approximately 99:1 (producer:non-producer). We incubated these in 5 ml iron limited medium (CAA) in the presence and absence of 50 *μ*M PQS for 24 h at 37°C with agitation at 200 rpm. To measure relative abundance of the strains, we plated the co-cultures before and after incubation, and counted total colonies and luminescent colonies. Relative fitness was calculated using the formula w = p_1_(1 − p_0_)/p_0_(1 − p_1_) where p_0_ and p_1_ are the proportion of non-producing mutants in the population before and after incubation respectively [28].

### (d) Statistical analyses

The effect of PQS and HHQ supplementation on growth, and siderophore production were all analyzed using the ordered heterogeneity approach [29]. This allows for the evaluation of an ordered alternative hypothesis but does not require the fitting of curves. We chose this approach because our question is about the effect of increasing concentrations of PQS but without any concern for the exact shape of these relationships. For each test, we calculated the test statistic r_s_P_c_ = r_S_ * (1 − *p*). p is the p-value from an ANOVA of raw data with concentration of PQS or HHQ fitted as a categorical variable, and r_S_ is the absolute value of the Spearman's rank correlation coefficient between the means of the relevant independent variable for each level of PQS or HHQ, and the concentration of PQS or HHQ. The relative fitness of a siderophore non-producer in iron limiting conditions, and the effect of PQS supplementation on the relative fitness, were examined using t-tests. All statistical analyses were performed using R 3.0.2 [30].

## 3. RESULTS

### (a) PQS-induced iron starvation increases the production of costly siderophores

To test whether PQS increased production of siderophores, we measured the amount of pyoverdine and pyochelin in cultures of a PQS-deficient mutant (PAO1Δ*pqsAH*) grown in LB, and supplemented with exogenous synthetic PQS at varying concentrations. We found that PQS reduced growth and increased the per-cell concentrations of pyoverdine and pyochelin in a concentration-dependent manner (figure 1a-c, ordered heterogeneity tests: growth F_6,28_ = 179, p < 0.001, r_s_ = − 0.964, r_s_P_c_ = 0.964, p < 0.05; pyoverdine F_6_,_28_ = 1311, p < 0.001, r_s_ = 0.643, r_s_P_c_ = 0.643, p < 0.05; pyochelin F_6_,_28_ = 2222, p < 0.001, r_s_ = 0.75, r_s_P_c_ = 0.750, p < 0.05). Since PQS plays a role in cell-cell communication, it is possible that reduced growth and induced iron scavenging are the result of QS-dependent regulation of gene expression. To exclude this possibility, we repeated the experiments in the presence of 2-heptyl-4-hydroxyquinoline (HHQ), the immediate precursor of PQS [31]. HHQ does not bind iron, but maintains a signalling role in cell-cell communication [22,31]. We found that increasing concentrations of HHQ did not significantly affect growth, pyochelin or pyoverdine production (figure 1d-f; ordered heterogeneity tests: growth F_6_,_28_ = 9.1, p < 0.001, r_s_ = 0.214, r_s_P_c_ = 0.214, p > 0.05; pyoverdine F_6_,_28_ = 6.6, p < 0.001, r_s_ = 0.25, r_s_P_c_ = 0.250, p > 0.05; pyochelin F_6_,_28_ = 24.4, p < 0.001, r_s_ = −0.071, r_s_P_c_ = 0.071, p > 0.05). Overall, we conclude that it is the iron chelating activity of PQS, and not its signal function, that triggers an iron starvation response: cells increase their production of iron scavenging siderophores and either a metabolic burden or slower uptake due to the presence of a chelator leads to poorer growth.

**Figure 1.**
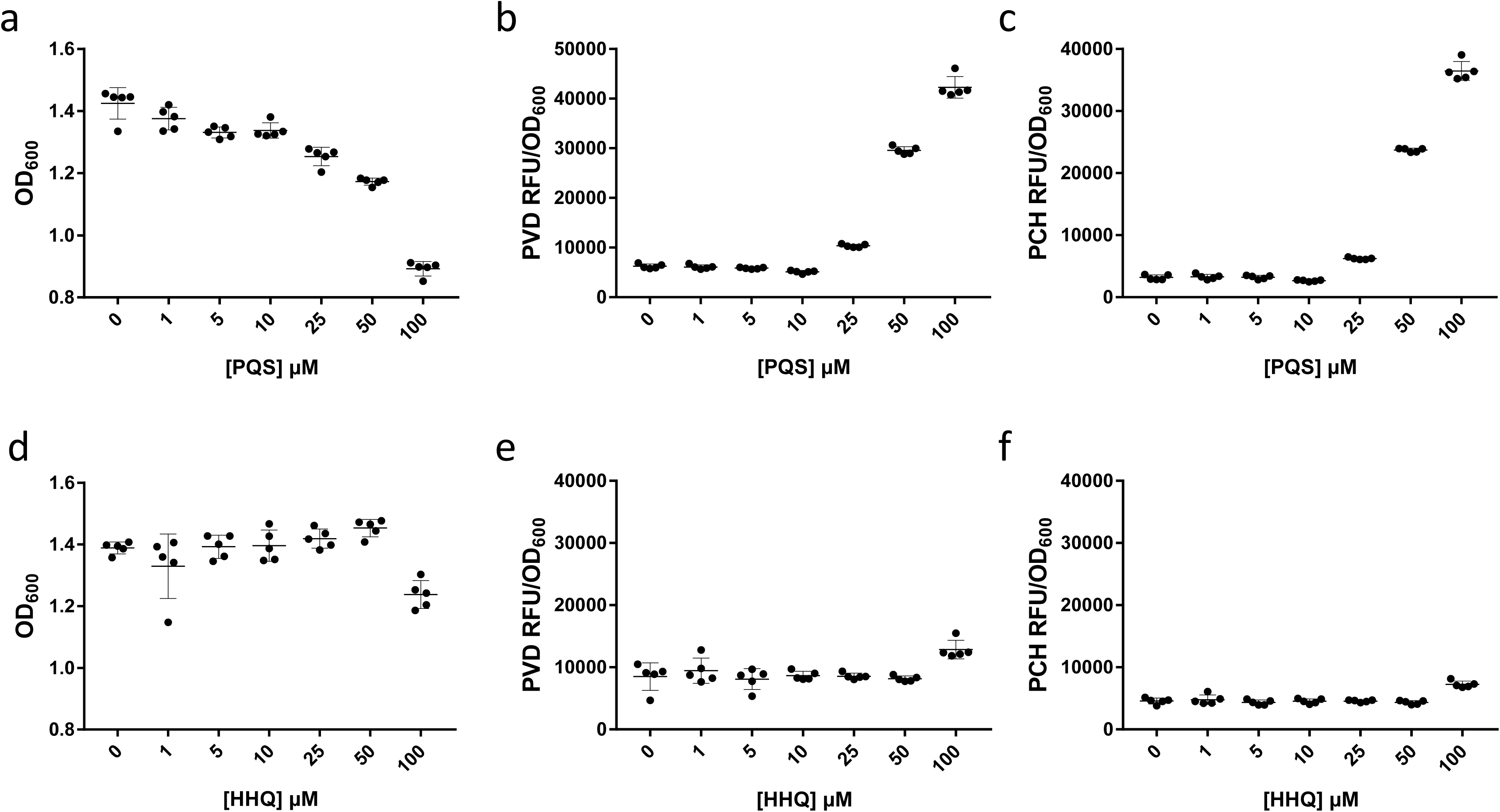
PQS causes iron starvation in *P. aeruginosa* cultures. (a) Increasing concentrations of exogenously added PQS decrease the growth of a PQS mutant (PAO1Δ*pqsAH*), in iron rich conditions and increase the production of the iron scavenging molecules (b) pyoverdine (PVD) and (c) pyochelin (PCH). (d-e) Iron starvation effects are not seen with the addition of HHQ, the biosynthetic precursor to PQS that does not bind iron. Error bars represent the standard deviation of 5 independent measurements.

### (b) PQS increases intra-specific competition for iron

The production of siderophores is a social trait that can be exploited by non-producing cheats [23]. We therefore predicted that intra-specific social competition over iron would intensify in the presence of exogenous PQS, due to the greater pool of siderophores available and the concomitant cost to producer growth. First we looked at the effect of PQS on monocultures of a PAO1 wild type and a strain defective in the production of both pyoverdine and pyochelin (PAO1Δ*pvdD/pchEF)* in iron limited media. The PAO1Δ*pvdD/pchEF* strain reached slightly higher optical densities than PAO1 in CAA media (p<0.001), but we found that PQS reduced the fitness of both strains which is consistent with the iron chelation effects of PQS (figure *2a*). We compared the PQS effect against the effect of transferrin, a chelator previously used in iron limited media siderophore experiments [25,32]. We found similar reductions in growth which shows that the PQS iron-chelating effect is comparable with that of transferrin (figure 2a). Consistent with existing work on the social dynamics of siderophore production, we found that the PAO1Δ*pvdD/pchEF* mutant functioned as a ‘cheat’ in iron-limiting conditions, having a relative fitness >1 when grown in co-culture with the wild type (figure *2b*; t_1_,_10_ = 3.32, *p* < 0.01), although the small increase in the fitness of the mutant (figure *2a*) could partially explain this finding. In line with our hypothesis, the addition of PQS significantly increased the relative fitness of the mutant (figure *2b*; F_1,10_ = 95.4, *p* < 0.001).

**Figure 2.**
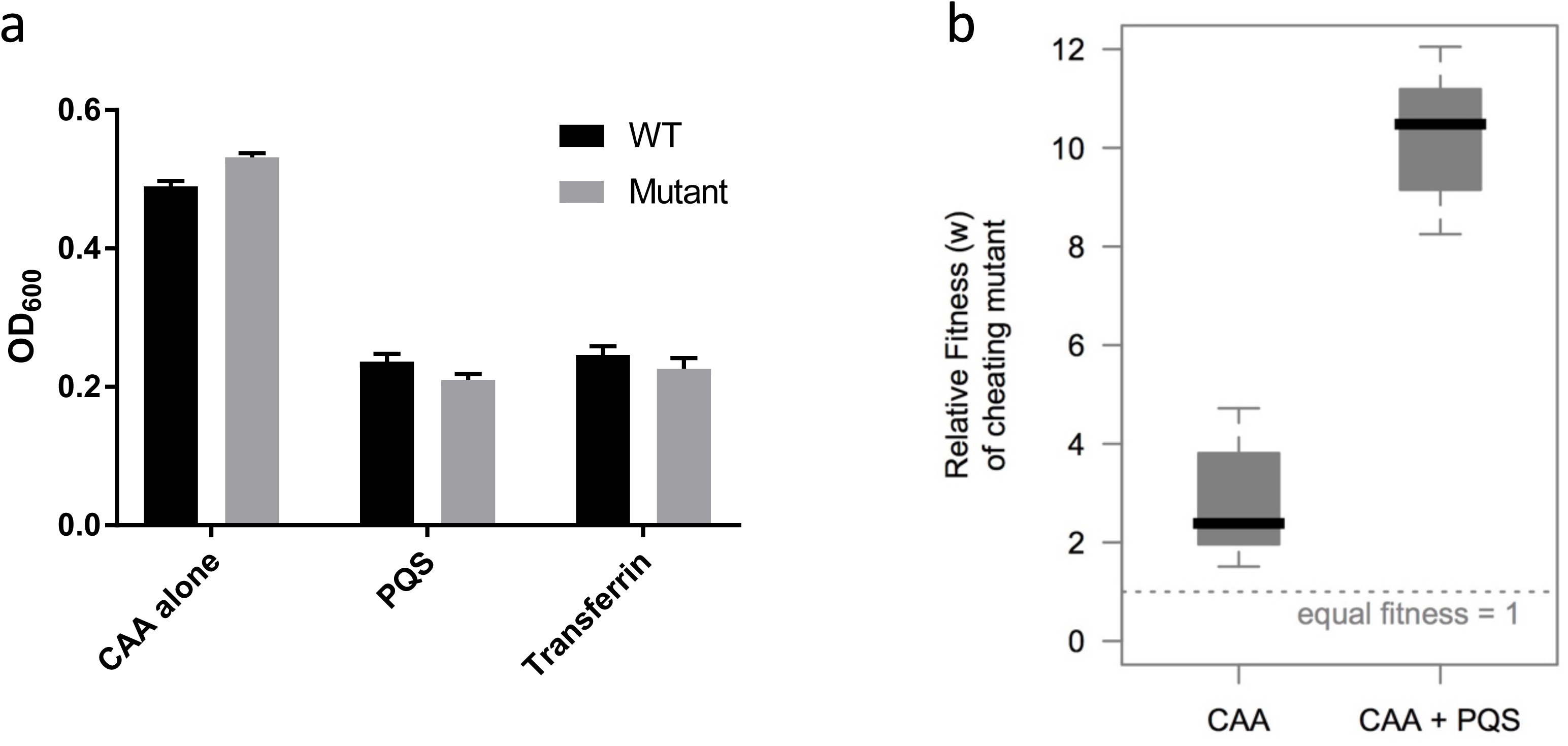
PQS increases the relative fitness of a siderophore cheat. (a) Monocultures of either PAO1 wild type (WT) or a double *pvdD/pchEF* mutant (Mutant) grown in CAA with either no supplementation or supplementation with either 50 *μ*M PQS or 100 *μ*g/ml transferrin. Error bars represent the standard deviation of 5 independent measurements. (b) A siderophore non-producing mutant gains a relative fitness advantage in co-culture with a siderophore producer in iron limiting conditions. When 50 *μ*M PQS is added to the culture this relative advantage increases due to increased siderophore output of the producer and subsequent increase in exploitation by the non-producer. The dashed line indicates the value of relative fitness (w = 1) at which both producer and non-producer have equal fitness. The box-plots indicate the median (red), the interquartile range (box) and the extreme values (whiskers).

## 4. DISCUSSION

Here we show, for the first time, that environmental modification via a QS molecule affects the selection for public goods that are not, as far as we are aware, directly regulated by QS. Specifically, we show that the iron-chelating properties of PQS lead to increased production of costly siderophores and consequently, increased relative fitness of a siderophore cheat. We found that the addition of synthetic PQS to cultures of *P. aeruginosa* results in a concentration-dependent decrease in bacterial fitness (growth) and an increase in the production of the siderophores pyoverdine and pyochelin (figure 1). The biosynthetic precursor of PQS, HHQ (which does not bind iron), had only a small effect on the production of pyoverdine but no effect on the production of pyochelin or on growth (figure 1).

We hypothesised that this effect of PQS would enhance the relative fitness payoff of siderophore non-producing cheats in competition with the wild type. Consistent with this hypothesis, we show that when siderophore production is increased by PQS in an iron-limited environment, this leads to an increase in the relative fitness of a cheating mutant (figure 2). The increased relative fitness of a cheat in the presence of PQS is likely due to a combination of the increased availability of siderophores to exploit, and the increased costs paid by siderophore producing cells. Our findings complement and build upon previous work, which showed that when less iron is available to cells, this results in greater production of siderophores, and an increase in the relative fitness of cheats [33]. In previous work, the authors artificially modified iron levels in the growth medium [33]. Our work differs in that we show that direct modification of iron levels in the environment by a QS molecule can alter selection for siderophore production.

Overall, our work builds upon a growing body of experimental studies exploring the complexities of cooperation in *P. aeruginosa*, an organism that is an excellent laboratory model for applying and testing and extending social evolution theory [1,2,6,8,11,13,34,35]. Microbes produce a diverse array of public goods, and little is known about how social traits interact with each other either directly or indirectly [36], although recent work has shown an interconnection between pyoverdine and pyochelin production [24]. Put another way, to what extent does the production of one social trait affect the social dynamics of another trait(s)? Existing examples include (a) the direct regulatory effect of communication on the production of public goods [7–9], and (b) the genetic linkage of traits via pleiotropy [37]. Future work in this area should continue to highlight and demonstrate which traits are social in microbes [38,39], but also begin to focus efforts on how apparently discrete traits interact, and how this affects population ecology and evolution within environments. This will require experiments that reveal the fitness effects of trait linkage, and also experiments to unravel the mechanisms by which traits are linked.

Given that we have shown PQS production enhances *P. aeruginosa* vulnerability to siderophore cheating, this suggests there are ecological and biological role(s) of PQS beyond its well documented role as a QS signal [16–18]. 2-alkyl-4(1*H*)-quinolones (including HHQ) have previously been shown to be produced by several bacterial species, but to date, PQS has only been shown to be produced by *P. aeruginosa* [18,40], suggesting that PQS may have evolved functions distinct from signalling. One possibility previously suggested, is that PQS-bound iron associates with the bacterial envelope making it easier for dedicated siderophores to shuttle iron into the cell. This could ensure that metabolically expensive siderophores are not easily lost to other cells [22]. Such a mechanism could help to reduce siderophore cheating, but our data shows that siderophore cheats flourish when PQS is present. Another role could be in ‘privatising’ iron for *P. aeruginosa* when it is in competition with other bacterial species. In this case, PQS-bound iron could reduce the availability of iron for heterospecific competitors whilst making it easier for pyoverdine and pyochelin to transport iron into cells. Future work focusing on understanding the ecology between bacterial species could help to unravel these interactions.

## DATA ACCESSIBILITY

Data and R code for analysing the data are available in Dryad: doi:10.5061/dryad.m13t1.

## AUTHOR CONTRIBUTION

R.P. and S.P.D. designed the experiments. R.P., F.H., A.C.S, and S.E. collected and analysed the data in the laboratory. R.P., F.H., L.M. and A.C.S. conducted the data analysis. R.P., L.M., P.W. and S.P.D wrote the paper. All authors revised and approved the manuscript.

## ETHICS STATEMENT

There are no ethical concerns to address in this paper.

## ACKNOWLEDGEMENTS

We thank Stuart West for comments on the manuscript.

## Bibliography

1. West, S. A., Griffin, A. S., Gardner, A. & Diggle, S. P. 2006 Social evolution theory for microorganisms. Nat Rev. Microbiol 4, 597–607. (doi:10.1038/nrmicro1461)

2. West, S. A., Diggle, S. P., Buckling, A., Gardner, A. & Griffins, A. S. 2007 The social lives of microbes. Annu Rev Ecol Evol S 38, 53–77. (doi:10.1146/Annurev.Ecolsys.38.091206.095740)

3. Hamilton, W. D. 1964 The genetical evolution of social behaviour. I. Journal of Theor Biol 7, 1–16.

4. Darch, S. E., West, S. A., Winzer, K. & Diggle, S. P. 2012 Density-dependent fitness benefits in quorum-sensing bacterial populations. Proc Natl Acad Sci USA 109, 8259–8263. (doi:10.1073/pnas.1118131109)

5. Schuster, M., Sexton, D. J., Diggle, S. P. & Greenberg, E. P. 2013 Acyl-homoserine lactone quorum sensing: from evolution to application. Ann Rev Microbiol 67, 43–63.(doi:10.1146/annurev-micro-092412-155635)

6. West, S. A., Winzer, K., Gardner, A. & Diggle, S. P. 2012 Quorum sensing and the confusion about diffusion. Trends Microbiol 20, 586–594. (doi:10.1016/j.tim.2012.09.004)

7. Brown, S. P. & Johnstone, R. A. 2001 Cooperation in the dark: signalling and collective action in quorum-sensing bacteria. Proc Biol Sci 268, 961–965.(doi:10.1098/rspb.2001.1609)

8. Diggle, S. P., Griffin, A. S., Campbell, G. S. & West, S. A. 2007 Cooperation and conflict in quorum-sensing bacterial populations. Nature 450, 411–414. (doi:10.1038/nature06279)

9. Czaran, T. & Hoekstra, R. F. 2009 Microbial communication, cooperation and cheating: quorum sensing drives the evolution of cooperation in bacteria. PLOS ONE 4, e6655. (doi:10.1371/journal.pone.0006655)

10. Rumbaugh, K. P., Diggle, S. P., Watters, C. M., Ross-Gillespie, A., Griffin, A. S. & West, S. A. 2009 Quorum sensing and the social evolution of bacterial virulence. Curr Biol 19, 341–345. (doi:10.1016/j.cub.2009.01.050)

11. Popat, R., Crusz, S. A., Messina, M., Williams, P., West, S. A. & Diggle, S. P. 2012 Quorum-sensing and cheating in bacterial biofilms. Proc Biol Sci 279, 4765–4771. (doi:10.1098/rspb.2012.1976)

12. Pollitt, E. J., West, S. A., Crusz, S. A., Burton-Chellew, M. N. & Diggle, S. P. 2014 Cooperation, quorum sensing, and evolution of virulence in *Staphylococcus aureus*. Infect Immun 82, 1045–1051. (doi:10.1128/IAI.01216-13)

13. Popat, R. et al. 2015 Conflict of interest and signal interference lead to the breakdown of honest signaling. Evolution 69, 2371–2383. (doi:10.1111/evo.12751)

14. Williams, P. & Camara, M. 2009 Quorum sensing and environmental adaptation in *Pseudomonas aeruginosa:* a tale of regulatory networks and multifunctional signal molecules. Curr Opin Microbiol 12, 182–191. (doi:10.1016/j.mib.2009.01.005)

15. Schertzer, J. W., Boulette, M. L. & Whiteley, M. 2009 More than a signal: non-signaling properties of quorum sensing molecules. Trends Microbiol 17, 189–195. (doi:10.1016/j.tim.2009.02.001)

16. Pesci, E. C., Milbank, J. B., Pearson, J. P., McKnight, S., Kende, A. S., Greenberg, E. P. & Iglewski, B. H. 1999 Quinolone signaling in the cell-to-cell communication system of *Pseudomonas aeruginosa*. Proc Natl Acad Sci USA 96, 11229–11234.

17. Dubern, J. F. & Diggle, S. P. 2008 Quorum sensing by 2-alkyl-4-quinolones in *Pseudomonas aeruginosa* and other bacterial species. Mol Biosystems 4, 882–888. (doi:10.1039/b803796p)

18. Heeb, S., Fletcher, M. P., Chhabra, S. R., Diggle, S. P., Williams, P. & Camara, M. 2011 Quinolones: from antibiotics to autoinducers. FEMS Micro Rev 35, 247–274. (doi:10.1111/j.1574-6976.2010.00247.x)

19. Mashburn, L. M. & Whiteley, M. 2005 Membrane vesicles traffic signals and facilitate group activities in a prokaryote. Nature 437, 422–425. (doi:10.1038/nature03925)

20. Haussler, S. & Becker, T. 2008 The Pseudomonas quinolone signal (PQS) balances life and death in *Pseudomonas aeruginosa* populations. PLOS Path 4, e1000166. (doi:10.1371/journal.ppat.1000166)

21. Bredenbruch, F., Geffers, R., Nimtz, M., Buer, J. & Haussler, S. 2006 The *Pseudomonas aeruginosa* quinolone signal (PQS) has an iron-chelating activity. Environ Microbiol 8, 1318–1329. (doi:10.1111/j.1462-2920.2006.01025.x)

22. Diggle, S. P. et al. 2007 The *Pseudomonas aeruginosa* 4-quinolone signal molecules HHQ and PQS play multifunctional roles in quorum sensing and iron entrapment. Chem Biol 14, 87–96. (doi: 10.1016/j.chembiol.2006.11.014)

23. Ratledge, C. & Dover, L. G. 2000 Iron metabolism in pathogenic bacteria. Ann Rev Microbiol 54, 881–941. (doi:10.1146/annurev.micro.54.1.881)

24. Ross-Gillespie, A., Dumas, Z. & Kummerli, R. 2015 Evolutionary dynamics of interlinked public goods traits: an experimental study of siderophore production i. Pseudomonas aeruginosa. J Evol Biol 28, 29–39. (doi:10.1111/jeb.12559)

25. Griffin, A. S., West, S. A. & Buckling, A. 2004 Cooperation and competition in pathogenic bacteria. Nature 430, 1024–1027. (doi:10.1038/nature02744)

26. Harrison, F., Browning, L. E., Vos, M. & Buckling, A. 2006 Cooperation and virulence in acute *Pseudomonas aeruginosa* infections. BMC Biol 4, 21. (doi:10.1186/1741-7007-4-21)

27. Dumas, Z., Ross-Gillespie, A. & Kummerli, R. 2013 Switching between apparently redundant iron-uptake mechanisms benefits bacteria in changeable environments. Proc Biol Soc 280, 20131055–20131055. (doi:10.1098/rspb.2013.1055)

28. Ross-Gillespie, A., Gardner, A., West, S. A. & Griffin, A. S. 2007 Frequency dependence and cooperation: theory and a test with bacteria. Am Nat 170, 331–342. (doi:10.1086/519860)

29. Rice, W. R. & Gaines, S. D. 1994 Extending nondirectional heterogeneity tests to evaluate simply ordered alternative hypotheses. Proc Natl Acad Sci USA 91, 225–226.

30. Ihaka, R. & Gentleman, R. 1996 R: a language for data analysis and graphics. J Comp Graph Stats 5, 299–314.

31. Deziel, E., Lepine, F., Milot, S., He, J., Mindrinos, M. N., Tompkins, R. G. & Rahme, L. G. 2004 Analysis of *Pseudomonas aeruginosa* 4-hydroxy-2-alkylquinolines (HAQs) reveals a role for 4-hydroxy-2-heptylquinoline in cell-to-cell communication. Proc Natl Acad Sci USA 101, 1339–1344. (doi:10.1073/pnas.0307694100)

32. Kummerli, R., Santorelli, L. A., Granato, E. T., Dumas, Z., Dobay, A., Griffin, A. S. & West, S. A. 2015 Co-evolutionary dynamics between public good producers and cheats in the bacterium *Pseudomonas aeruginosa*. J Evol Biol 28, 2264–2274. (doi:10.1111/jeb.12751)

33. Kummerli, R., Jiricny, N., Clarke, L. S., West, S. A. & Griffin, A. S. 2009 Phenotypic plasticity of a cooperative behaviour in bacteria. J Evol Biol 22, 589–598. (doi:10.1111/j.1420-9101.2008.01666.x)

34. Rumbaugh, K. P., Griswold, J. A., Iglewski, B. H. & Hamood, A. N. 1999 Contribution of quorum sensing to the virulence of *Pseudomonas aeruginosa* in burn wound infections. Infect Immun 67, 5854–5862.

35. Sandoz, K. M., Mitzimberg, S. M. & Schuster, M. 2007 Social cheating in *Pseudomonas aeruginosa* quorum sensing. Proc Natl Acad Sci USA 104, 15876–15881. (doi:10.1073/pnas.0705653104)

36. Brown, S. P. & Taylor, P. D. 2010 Joint evolution of multiple social traits: a kin selection analysis. Proc Biol Soc 277, 415–422. (doi:10.1098/rspb.2009.1480)

37. Harrison, F. & Buckling, A. 2009 Siderophore production and biofilm formation as linked social traits. ISME J 3, 632–634. (doi:10.1038/ismej.2009.9)

38. Ghoul, M., West, S. A., Diggle, S. P. & Griffin, A. S. 2014 An experimental test of whether cheating is context dependent. J Evol Biol 27, 551–556. (doi:10.1111/jeb.12319)

39. Ghoul, M., Griffin, A. S. & West, S. A. 2014 Toward an evolutionary definition of cheating. Evolution 68, 318–331. (doi:10.1111/evo.12266)

40. Diggle, S. P., Lumjiaktase, P., Dipilato, F., Winzer, K., Kunakorn, M., Barrett, D. A., Chhabra, S. R., Camara, M. & Williams, P. 2006 Functional genetic analysis reveals a 2- Alkyl-4-quinolone signaling system in the human pathogen *Burkholderia pseudomallei* and related bacteria. Chem Biol 13, 701–710. (doi:10.1016/j.chembiol.2006.05.006)

